# Novel genetic sex markers reveal unexpected lack of, and similar susceptibility to, sex reversal in free-living common toads in both natural and anthropogenic habitats

**DOI:** 10.1101/2021.10.18.464681

**Authors:** Edina Nemesházi, Gábor Sramkó, Levente Laczkó, Emese Balogh, Lajos Szatmári, Nóra Vili, Nikolett Ujhegyi, Bálint Üveges, Veronika Bókony

**Affiliations:** Conservation Genetics Research Group, Department of Ecology, University of Veterinary Medicine Budapest, István u. 2, 1078 Budapest, Hungary; Lendület Evolutionary Ecology Research Group, Plant Protection Institute, Centre for Agricultural Research, Eötvös Loránd Research Network, Herman Ottó u. 15, 1022 Budapest, Hungary; MTA-DE Lendület Evolutionary Phylogenomics Research Group, Egyetem tér 1, 4032 Debrecen, Hungary; Molecular Ecology and Evolution, School of Natural Sciences, Bangor University, Bangor LL57 2UW, Wales, United Kingdom

**Keywords:** amphibians, molecular sex markers, sex change, sex-chromosome identification, human-induced environmental change, feminization

## Abstract

Anthropogenic environmental changes are affecting biodiversity and microevolution worldwide. Ectothermic vertebrates are especially vulnerable, since their sexual development can be disrupted by environmental changes, which can cause sex reversal, a mismatch between genetic and phenotypic sex, potentially leading to sex-ratio distortion and population decline. Despite these implications, we have scarce empirical knowledge on the incidence of sex reversal in nature. Populations in anthropogenic environments may experience sex reversal more frequently, or alternatively, they may adapt to resist sex reversal. To test these alternative hypotheses, we developed PCR-based genetic sex markers for the common toad *(Bufo bufo)*. We assessed the prevalence of sex reversal in wild populations living in natural, agricultural and urban habitats, and the susceptibility of the same populations to two ubiquitous estrogenic pollutants in a common-garden experiment. We found negligible sex-reversal frequency in free-living adults despite the presence of various endocrine-disrupting pollutants in their breeding ponds. Individuals from different habitat types showed similar susceptibility to sex reversal in the laboratory: all genetic males developed female phenotype when exposed to 1 µg/L 17α- ethinylestradiol (EE2) during larval development, whereas no sex reversal occurred in response to 1 ng/L EE2 and a glyphosate-based herbicide with 3 µg/L or 3 mg/L glyphosate. The latter results do not support that populations in anthropogenic habitats would have either increased propensity for or higher tolerance to chemically induced sex reversal. Thus, the surprisingly low sex-reversal frequency in wild toads compared to other ectothermic vertebrates studied before might indicate idiosyncratic, potentially species-specific resistance to sex reversal.

## Introduction

Anthropogenic environmental change confronts wildlife with challenges that call for rapid phenotypic or genetic adaptations. For example, both urban and agricultural land use loads the environment with various chemical pollutants including pesticides, heavy metals, road de-icers, pharmaceuticals and industrial products, while climate change increases the frequency and intensity of heat waves and other extreme weather events. Understanding how these human-induced environmental changes affect the ecosystem, and whether and how wildlife can adapt to overcome these challenges, is an important mission of current evolutionary ecology (Tilman et al., 2017).

In ectothermic animals, various environmental stimuli can cause a developmental effect rarely seen in endotherms: sex reversal, whereby individuals exposed to such stimuli during their embryonic or larval life-phase develop the sexual phenotype opposite to their genetic sex (Baroiller & D’Cotta, 2016; Flament, 2016; Whiteley, Castelli, Dissanayake, Holleley, & Georges, 2021). Sex reversal occurs in fish, amphibians, and reptiles in nature (Alho, Matsuba, & Merilä, 2010; Baroiller & D’Cotta, 2016; Lambert, Tran, Kilian, Ezaz, & Skelly, 2019; Nemesházi et al., 2020; Whiteley et al., 2021; Xu et al., 2021), and theoretical studies caution that it may have far-reaching consequences including skewed sex ratios, sex-chromosome evolution, and even population extinction (Bókony, Kövér, Nemesházi, Liker, & Székely, 2017; Grossen, Neuenschwander, & Perrin, 2011; Nemesházi, Kövér, & Bókony, 2021; Perrin, 2009; Schwanz, Georges, Holleley, & Sarre, 2020; Wedekind, 2017). Laboratory experiments show that sex reversal can be induced by anthropogenic stressors like chemical pollution and elevated temperature (Flament, 2016; Lambert, Smylie, Roman, Freidenburg, & Skelly, 2018; Mikó et al., 2021; Tamschick et al., 2016), thus, we may expect that the contemporary and future increase in the levels of anthropogenic stressors will influence the rates of sex reversal in free-living populations of ectothermic vertebrates. This influence is conceivable in at least two ways.

On the one hand, sex reversal may happen more often in areas where the chemical and thermal stimuli triggering it are more pervasive, such as in agricultural areas polluted by pesticides and in urban heat islands. This may be simply a consequence of sex-reversing stimuli being more frequent in such habitats, but alternatively or additionally, frequent sex reversal might also be an adaptive response to anthropogenic environments. Adjusting phenotypic sex to environmental conditions can be adaptive (Geffroy & Douhard, 2019), and sex reversal might be an evolved mechanism for achieving the sexual phenotype that best matches the environment, similarly to environmental sex determination (Schwanz & Georges, 2021). If sex reversal in anthropogenic environments is adaptive, populations persisting in anthropogenic habitats may evolve higher propensity for sex reversal. This might be facilitated by genetic and epigenetic inheritance of the propensity to develop into one sex or the other (McGaugh & Janzen, 2011; Piferrer & Anastasiadi, 2021).

On the other hand, sex reversal may be costly in terms of fitness. For example, reproductive performance of fish is reduced by sex reversal (Pandian & Sheela, 1995; Alistair McNair Senior, Nat Lim, & Nakagawa, 2012), as well as by intersex, a form of imperfect sex reversal when the gonads contain both male and female tissues (Fuzzen, Bennett, Tetreault, McMaster, & Servos, 2015; Harris et al., 2011). Due to their sex-chromosome genotype, sex-reversed individuals may be unable to produce daughters or sons (Wedekind, 2017), and therefore may be selected against by sex-ratio selection (Schwanz & Georges, 2021; see also Fig S11a,d in Nemesházi *et al*., 2021a). Also, sex-reversed individuals may perform poorly in traits that influence survival (Mikó et al., 2021; Nemesházi et al., 2020) or sexually selected traits (Nemesházi, Kövér, et al., 2021). In such situations, we can expect resistance to sex reversal to be adaptive in environments where sex-reversing stressors are pervasive. As a result of such adaptation, populations exposed to sex-reversing environments might maintain the same frequency of sex reversal as unexposed populations.

Putting these ideas to the test empirically has been hindered by the difficulty of diagnosing sex reversal in non-model organisms. Due to the high evolutionary lability and homomorphy of sex chromosomes in ectothermic vertebrates, genetic sexing methods are available only for a handful of species (e.g. Alho *et al*., 2010; Baroiller & D’Cotta, 2016; Tamschick *et al*., 2016; Lambert *et al*., 2019; Nemesházi *et al*., 2020; Whiteley *et al*., 2021; Xu *et al*., 2021). In two of those species, recently developed genetic sex markers have been used to investigate whether sex reversal is more prevalent in anthropogenic habitats, and they reported contradictory answers: yes in one frog species (Nemesházi et al., 2020) but no in another (Lambert et al., 2019). Furthermore, no study, to our knowledge, has yet tested whether animal populations living in anthropogenic habitats have increased or reduced inherent propensity for sex reversal.

In this study, we first aimed to produce a reliable molecular marker set for diagnosing genetic sex in the common toad (*Bufo bufo*), an anuran amphibian widespread in Eurasia that occupies a wide range of habitats from pristine woodlands to anthropogenic areas (Agasyan et al., 2009). This species has a female-heterogametic (ZZ/ZW) sex-chromosome system (Dufresnes et al., 2020), and is liable to chemically induced sex reversal (Hayes, 1998). Then, using our novel marker set, we investigated whether the frequency of sex reversal in toads differed between natural, agricultural, and urban habitats. Finally, we performed a common garden experiment to test whether toads originating from these three types of habitat differ in their susceptibility to sex reversal induced by chemical pollutants.

We focused on the sex-reversing effects of two endocrine disrupting chemical (EDC) compounds with high prevalence in surface water in agricultural and urban areas, respectively: glyphosate, the most used herbicide worldwide (Brovini et al., 2021), and 17α-ethinylestradiol (EE2), a common ingredient of contraceptives that pollutes natural water bodies *via* wastewater (Bhandari et al., 2015). Both EDCs may cause male-to-female sex reversal based on their effects on estrogenic enzymatic activities, female-skewed sex ratios, and intersex gonads (Bhandari et al., 2015; Howe et al., 2004; Lanctôt et al., 2014; Tamschick et al., 2016). As both chemicals have been in use for about half a century, we can expect resistance to have potentially evolved in populations chronically exposed to these pollutants. Similar, rapid evolutionary changes due to anthropogenic habitat alterations have been documented in various taxa, including evolved tolerance to lethal effects of pollutants (Brans, Almeida, & Fajgenblat, 2021; Cothran, Brown, & Relyea, 2013; Johnson & Munshi-South, 2017; Marques da Cunha, Maitre, & Wedekind, 2019; Reid et al., 2016). Here we test for altered susceptibility to sex reversal (a sub-lethal EDC effect) in common toad populations living in anthropogenic habitats.

## Methods

### Data collection

We captured 352 adult toads during the spawning seasons of 2016 and 2017 at 14 breeding sites in North-Central Hungary, which represented three habitat types: natural, agricultural, and urban areas, with 4–5 sites per habitat type (Table S1, Figure S1, Figure S2). We identified the phenotypic sex of adults by sexual characteristics: nuptial pads in males (N=216) and presence of eggs in females (N=136). We took a DNA sample from each individual (buccal swab or tissue sample) and stored it in 96% ethanol.

In 2017, we transferred 89 pairs of the captured adults to captivity and allowed them to spawn there, as described in an earlier paper (Bókony et al., 2018). Depending on the availability of females and their willingness to spawn in captivity, we had 1–15 egg strings from each of 11 sites out of the 14 sites sampled for adult DNA (36, 16, and 37 families from natural, agricultural, and urban sites, respectively; Table S1). When the tadpoles hatching from the captive-laid eggs reached the free-swimming stage (developmental stage 25; Gosner, 1960), we haphazardly selected six individuals from each family, distributed them among six treatments (control, solvent control, and four EDC treatments; N=534: Table S2; see below), and raised them for ca. 5 months after metamorphosis as described in an earlier paper (Ujhegyi & Bókony, 2020). Because the methodological details of these procedures have already been published, we repeat them in the Supplementary Information (sections 1 and 2) of the present paper, and only the most important aspects of the experiment are described below.

The control group was kept in clean water, and served as control for the glyphosate treatments, in which a glyphosate-based herbicide formulation (Glyphogan® Classic; Monsanto Europe S.A., Brussels, Belgium; containing 41.5 w/w% glyphosate and 15.5 w/w% polyethoxylated tallow amines) was added to the rearing water to maintain a nominal concentration of either 3 µg/L or 3 mg/L glyphosate. The solvent-control group, in which the rearing water contained 1 µL/L ethanol, served as control for the EE2 treatments, in which the nominal concentration was either 1 ng/L or 1 µg/L EE2, obtained by dissolving EE2 powder (Sigma E4876) in 96% ethanol and adding 1 µL of this solution to each litre of rearing water. Actual EDC concentrations were close to the nominal concentrations (Ujhegyi & Bókony, 2020). Both EDCs are documented to occur in our actual study ponds (Bókony et al., 2018). The lower and higher concentrations we used for each EDC represent the typical and maximum concentrations, respectively, detected in surface waters (Avar et al., 2016; Bhandari et al., 2015; Bókony et al., 2018; Brovini et al., 2021). The treatments lasted throughout the entire larval period for each individual, and were renewed twice a week at each water change.

When the toadlets (N=417) reached the age by which their gonads are completely differentiated (Ogielska & Kotusz, 2004), we euthanized them using MS-222, and identified whether each individual had testes or ovaries by dissection. We stored the gonads in 10% buffered formalin and later examined them histologically (for detailed methods, see Nemesházi *et al*., 2020). In a few cases when we could not unambiguously categorize the gonads as testes or ovaries based on gross anatomy and histology, we treated the phenotypic sex as uncertain. We stored the body of dissected toadlets in 96% ethanol until extracting DNA from a foot sample (see section 3 in Supplementary Information). Metadata for both adults and juveniles are publicly available on FigShare (Nemesházi, Sramkó, et al., 2021).

### Marker development and validation

For developing and validating genetic sex markers, we used a reduced-representation genomic library approach on toadlets with known phenotypic sex from the control group of the common garden experiment. Since these animals had not been exposed to any stimuli that are expected to cause sex reversal, they are likely to have phenotypic sex concordant with their genetic sex. First, we selected 24 non-sibling individuals sexed by gonad morphology (11 males, 13 females), representing 10 out of the 11 capture sites each by 1–4 toadlets. After DNA extraction (see section 3 in Supplementary Information), we applied Restriction-site Associated DNA Sequencing (RADseq) to identify sex-specific markers using the approach of Feron *et al*. (2021), which statistically examines RAD-tags for being significantly associated with *a priori* sex.

To generate RADseq data, we adopted the original RADseq protocol of Baird *et al*. (2008) and used dual-barcoded modified Illumina adapters. Next generation sequencing libraries were prepared using 210 ng gDNA as starting material from each isolate that was digested with the rare cutter restriction enzyme SbfI-HF (New England Biolabs, MA, USA). Custom P1 adapters were ligated at sticky ends, then isolates with a different adapter were pooled and sheared using a Bioruptor Pico machine (5 cycles of 30 s ‘on’ 30 s ‘off’). Libraries were size selected using the SPRI Select Kit (Beckman Coulter, CA, USA) to contain 300–600 base-pair (bp) long fragments only. Custom P2 adapters were ligated to the fragments and sub-libraries were pooled equimolarly. PCR enrichment used the Phusion High-Fidelity PCR Master Mix (New England Biolabs, MA, USA) and a decreased number of PCR cycles (14) compared to the original protocol of Baird *et al*. (2008). After a final size selection the quality and quantity of the library was checked on a Bioanalyzer (Agilent Technologies, CA, USA) device, then the library was sequenced on an Illumina HiSeq platform with 150 bp paired-end sequencing option at a commercially available service provider (Novogene Co. Ltd., Beijing, China).

Raw Illumina reads were demultiplexed and filtered using *process_radtags* from the Stacks v.2.2 pipeline (Rochette, Rivera-Colón, & Catchen, 2019). Adapter content was additionally checked and removed by using fastp v.0.20.1 (Chen, Zhou, Chen, & Gu, 2018). Using the ‘forward’ (R1) reads only, we screened the dataset for sex-specific reads of the 24 samples with known sex by RADSex v.1.1.2 (Feron et al., 2021). We identified significantly sex-linked markers by setting the significance threshold to the False Discovery Rate adjustment (Benjamini, Drai, Elmer, Kafkafi, & Golani, 2001) of the P=0.05 threshold adjusted for the number of tests (N=168 combinations of number of males and number of females in which the given marker is present; sex linkage tested by Pearson’s χ^2^ test of independence with Yates’ correction for continuity). We set the minimum read depth to one, thus allowing the discovery of the maximum number of potentially sex-linked markers. Read depth and distribution of sex-specific markers were checked with the *sgtr* package (Feron et al., 2021) in R (R Core Team, 2014). Since RADSex can only process reads that totally overlap (practically restricting this step to the ‘forward’ reads), we retained sequence information of the ‘reverse’ (R2) reads by picking the reads identified by RADSex as significant markers and assembling ‘contigs’ using the corresponding paired-end reads by using the GNU/Linux utility ‘grep’. First, we extracted all exact sequence matches of the forward reads to the output sequences of *radsex signif*, then searched for the read pairs by their unique read identifiers. These short reads were clustered into contigs by CD-HIT v.4.8.1 (Fu, Niu, Zhu, Wu, & Li, 2012) with a sequence similarity of 1.0 and PEAR v.0.9.6 (Zhang, Kobert, Flouri, & Stamatakis, 2014), as implemented in dDocent v.2.7.8 (Puritz, Hollenbeck, & Gold, 2014), that is designed for the *de novo* processing of RADseq datasets.

Using the NCBI Primer designing tool (https://www.ncbi.nlm.nih.gov/tools/primer-blast), we designed sequencing primers for polymerase chain reaction (PCR) amplification of potentially sex-linked loci from the assembled RAD loci in order to obtain Sanger sequences of them. After PCR-optimization (conditions for each sequencing primer pair are available in Table S3), loci that gave a bright PCR-product band in the expected size ranges on a 2% agarose gel were further processed. Such PCR products of three female and one male laboratory-raised juvenile toads from the control group were cut and purified from the gel using the NucleoSpin Gel and PCR Clean-up Kit (Macherey-Nagel) and were sequenced on a 3130xl Genetic Analyzer (Thermo Fisher Scientific) by a commercially available service provider (BIOMI Kft., Gödöllő, Hungary). We initially sequenced more females than males to obtain multiple copies from both sex chromosomes (i.e. a total of five Z and three W copies from the three ZW females and one ZZ male). Sequences from each locus were manually checked using the Staden software package (Bonfield, Smith, & Staden, 1995) and subsequently aligned in Mega (version 7.0.26). We focused on those loci where we both obtained unambiguous sequences and found sex-linked sequence differences, which were either single nucleotide polymorphisms (SNPs) or insertions and deletions (InDels). These loci were sequenced in 5 males and 5 females in total for the purpose of designing diagnostic sexing primers. For 4 markers, we developed optimized sexing primers as described in Supplementary Information section 4. When a Z/W InDel difference was large enough for detection on agarose gel, we designed sexing primers that would bind to both sex chromosomes (Z/W primers) and yield fragments of different length. For other loci, we added a digestion step where the W product was cut into two fragments by a restriction enzyme to be sex-specific. When the above methods failed, we included a third, W-specific primer in the sexing PCRs to obtain a clearly distinguishable W product along with the products of the Z/W sexing primers.

We subsequently tested the developed sexing method for each sex marker (c2, c5, c12 and c16; see sexing primers in Table 1) in 46 males and 36 females with unambiguous sexual phenotype from the control group, all being non-siblings and representing all 11 study sites that were used for the common garden experiment (including those used for sequencing and primer design). Additionally, we searched the common toad genome (NCBI GeneBank identifier: aBufBuf1.1) in order to identify the sex chromosomes based on our sex markers (see Supplementary Information section 5).

**Table 1.**
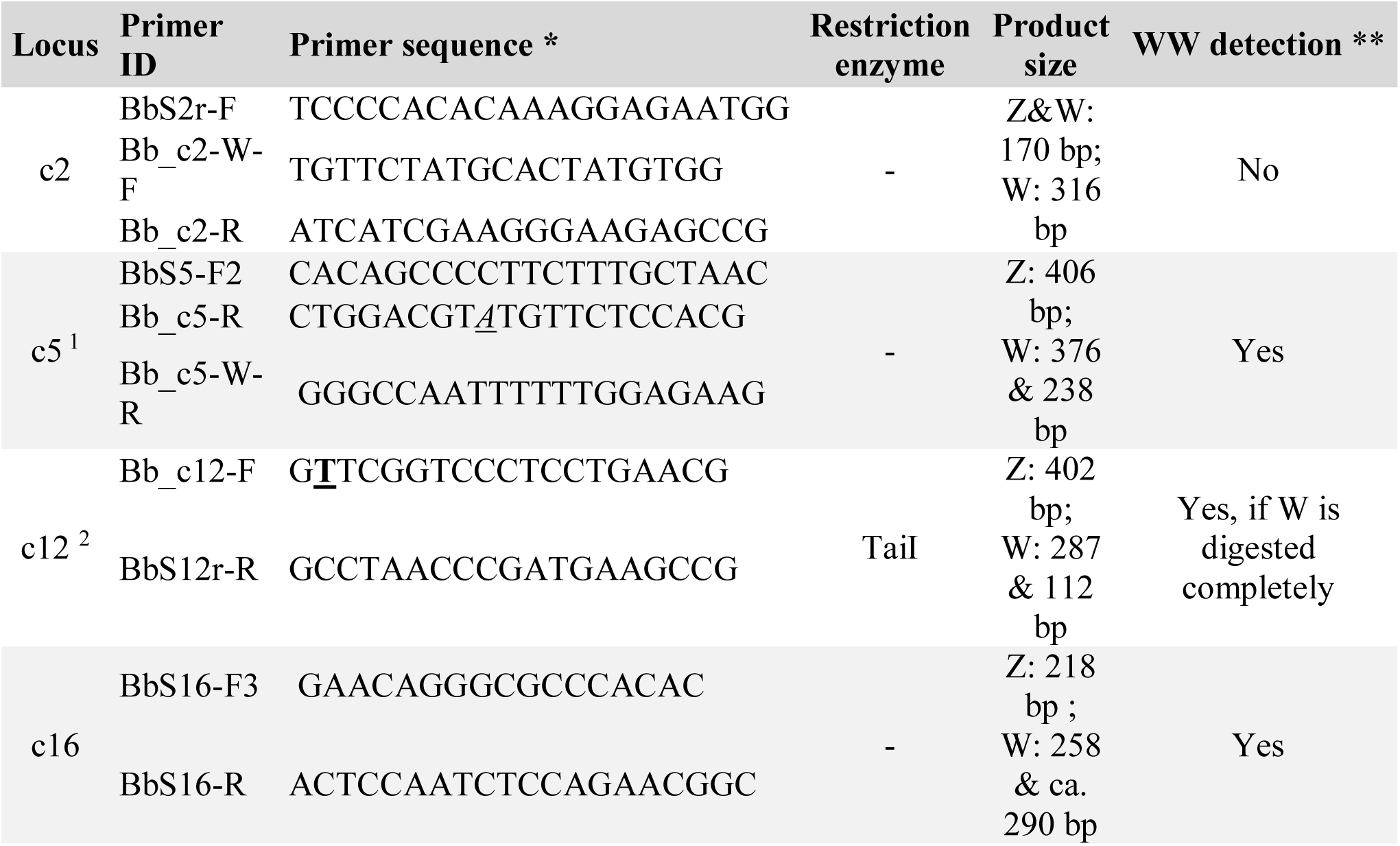
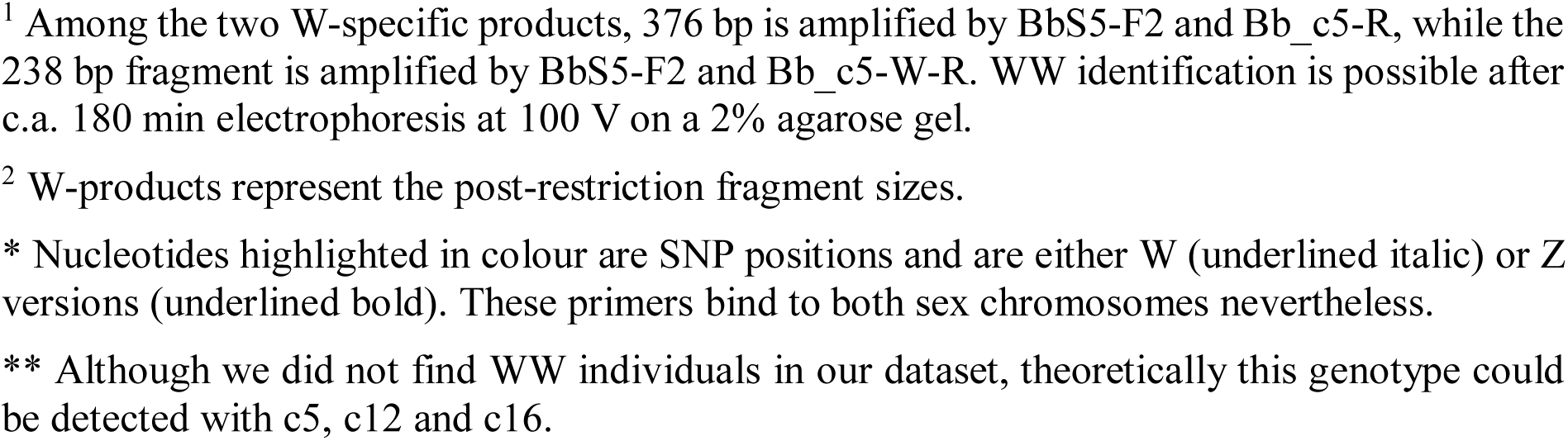
Sexing primers.

### Identification of sex reversals

We used our novel sexing primers to distinguish sex-reversed and sex-concordant individuals both among free-living adults and their juvenile offspring raised in our common-garden experiment. Preparation of DNA samples for subsequent analyses is described in section 3 in Supplementary Information. Sexing PCRs were performed on a Life ECO TC-96/G/H(b)C (Bioer) or a Biometra Tone 96G (Analytic Jena) instrument using one of the following two touch-down protocols. PCRs of c5, c12 and c16 were performed as follows: 2 min denaturation at 94°C followed by 13 cycles of 30 sec denaturation at 94°C, 30 sec annealing gradually decreasing from 68 to 64°C (−0.3°C per cycle) and 30 sec elongation at 72°C, followed by 22 more cycles with the same settings but constant 64°C annealing temperature, and a final 10 min extension step at 72°C. Amplification of c2 differed from the above PCR profile: the annealing temperature decreased from 66 to 60°C in 3 cycles (touch-down phase), followed by 32 cycles with an annealing temperature of 60°C. PCRs of all markers were performed in a total of 10 µl volume containing 2 µl FIREPol Master Mix (5x, ready to load; Solis BioDyne), 5 µl unquantified DNA for swab samples or 1 µl unquantified DNA for tissue samples, and varying amount of PCR primers and nuclease-free water. In case of the two-primer PCRs (c12 and c16), 0.4 pmol of each PCR primer (i.e. one forward and one reverse) was used. In case of the three-primer PCRs (c2 and c5), a mixture of 0.05 pmol BbS2r-F, 0.45 pmol of Bb_c2-W-F and 0.4 pmol of Bb_c2-R performed best for c2, while 0.8 pmol of BbS5-F2, 0.175 pmol of Bb_c5-W-R and 0.15 pmol of Bb_c5-R gave the best result for c5. In order to perform W-specific digestion for c12, we subsequently added 1.43 µl Tango Buffer (10x, Thermo Scientific), 0.72 µl TaiI restriction enzyme (10 U/µl, Thermo Scientific) and 2.85 µl nuclease-free water to the PCR product, resulting in a 15 µl final volume. Digestion was performed at 65°C for two hours. With each marker, genetic sex was identified by electrophoresis on 2% agarose (peqGOLD Electran, VWR Peqlab) gel (0.5% TBE buffer Thermo Scientific) stained with ECO Safe (Pacific Image Electronics Co., Ltd.).

Because both the Z-linked and W-linked PCR products of c16 were similarly bright on the agarose gel, and genotyping required only a simple PCR with one sexing-primer pair (i.e. no third primer or enzymatic restriction was necessary), we decided to use this marker for screening all those individuals from the common garden experiment that had not been sexed during the marker testing phase as well as the wild-caught adults. If the c16 genotype of an individual did not match its phenotype, we genotyped the individual for c12 as well, to ensure correct assignment of the genetic sex. Individuals with uncertain sexual phenotype were sexed for at least two additional markers besides c16.

## Results

### Marker development and validation

Based on 24 individuals, RADSex identified 17 significantly sex-linked markers (Figure 1), out of which 13 showed female-biased pattern (as expected under ZW/ZZ sex determination), but two of the latter were suspected to be paralogue sequences. During marker development, we concentrated on the remaining 11 RAD loci and designed sequencing primer pairs for each. Out of these, 9 primer pairs produced bright PCR products of the expected fragment size, and we obtained unambiguous sequences from 7 loci (Table S3). We found sex-linked InDel or SNP differences in the sequences of 4 loci hereafter referred to as c2, c5, c12 and c16 sex markers (sequences were registered at NCBI GenBank under the following accession numbers: OK507208-OK507215). NCBI genome blast search (see also Supplementary Information section 5) indicated that c12 was located on chromosome 5, whereas c2 and c16 were localized on two different, unplaced scaffolds: the former is currently suggested to belong to chromosome 6, but no such information is available on the latter. For c5, genome blast showed highly similar sequences on several different chromosomes, including multiple locations on chromosome 5.

**Figure 1.**
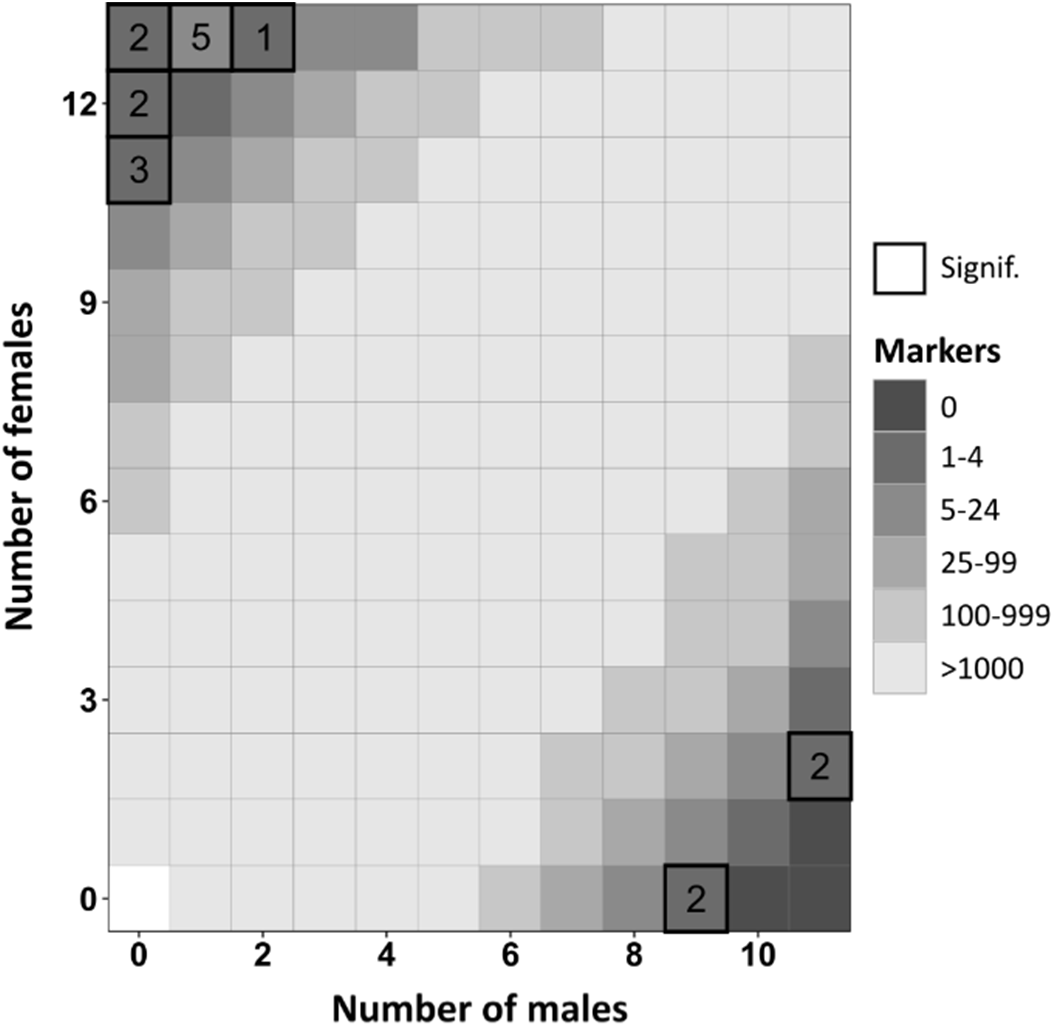
Tile plot showing the number of RADSex markers found in different sexes. Shade of each tile refers to the number of markers that were found in a given number of males and females. Thick black frames around tiles show significantly sex-linked markers, and the numbers within these tiles indicate the exact number of markers.

Final sexing PCR primers for our 4 sex-linked markers, product sizes and furtherdetails are shown in Table 1. Sexual genotypes based on each marker matched the sexual phenotype in all 82 non-sibling individuals chosen for marker validation, yielding 100% reliability for sexing with each of our four, newly devised markers.

### Identification of sex reversals

We successfully genotyped 349 wild-caught adults from 14 breeding ponds, while PCRs failed in three individuals (Table S1). We found 135 concordant ZW females and 213 concordant ZZ males, and a single sex-reversed individual, a phenotypic male from an agricultural site (‘Határrét’; see Figure S1) which was diagnosed as genetic female (ZW) by 3 out of 4 markers (c16, c2, and c12), while repeated PCRs with marker c5 gave ambiguous results.

We successfully genotyped all 417 toadlets that survived until phenotypic sexing in the common garden experiment. We detected no sex reversal in any of the treatment groups, except for the higher concentration of EE2 (Figure 2). In the latter treatment, all genetic males developed into phenotypic females, regardless of their original habitat type (Figure 2). The ovaries of the male-to-female sex-reversed individuals (ZZ females) were anatomically and histologically indistinguishable from the ovaries of concordant females (Figure S3). Additionally, 6 individuals with uncertain sexual phenotype were found to be genetically males (ZZ). Four of these toadlets, all originating from urban ponds, had intersex gonads (Figure S4); three of them had been treated with the lower concentration of the glyphosate-based herbicide and one with the lower concentration of EE2 (Figure 2). In the remaining two toadlets we could not unambiguously ascertain if the gonads were intersex or normal testes (Figure S4).

**Figure 2.**
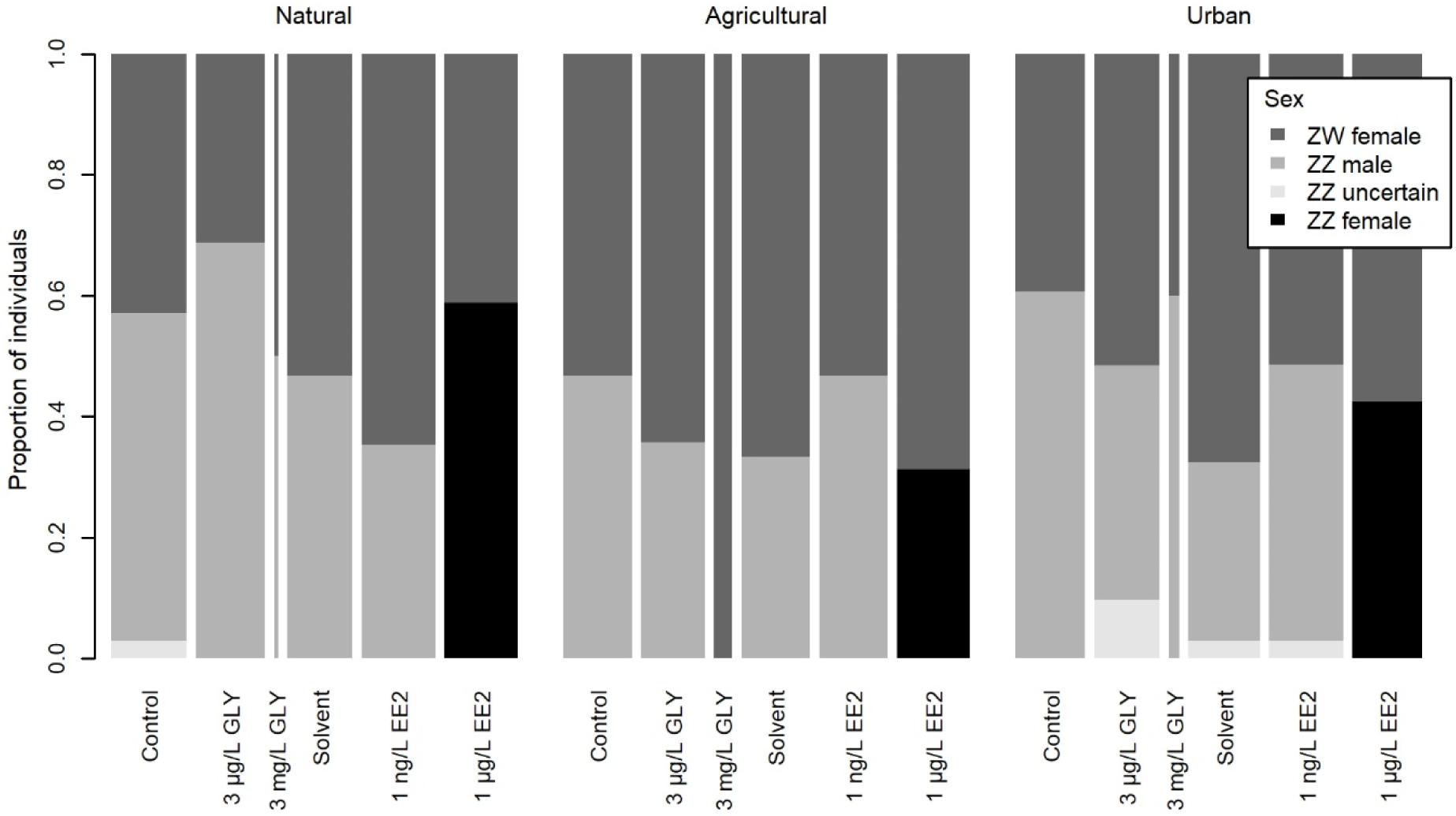
Proportion of concordant and sex-reversed individuals in each treatment group by habitat type of the parents’ capture site. Bar widths are proportional to sample size, which varied between 2 and 35 due to differences in survival (see Table S2).

## Discussion

The novel sex markers developed in our study confirmed that Hungarian populations of the common toad are female heterogametic, echoing recent findings from Switzerland (Dufresnes et al., 2020). Identification of the common toad’s sex chromosomes remained unresolved so far (Dufresnes et al., 2020). Genome BLAST showed that at least three of the four new sex markers are located on different scaffolds, but specific position of these scaffolds is unknown, except for one which is located on chromosome 5. One of the unplaced scaffolds however, was suggested to belong to chromosome 6. Some anuran species feature complex sex determination that can include multiple sex chromosomes (Gazoni et al., 2018; Roco et al., 2015). In such cases, sex chromosomes can form a meiotic chain, therefore multiple loci of different chromosomes may be inherited in linkage. Because each of our sex markers showed 100% sex linkage, we are confident that they are all located on the sex chromosome(s). Based on the available data, we propose chromosome pair 5 (most similar to chromosome 6 in *Xenopus tropicalis*) as candidate sex chromosome pair in the common toad, but the exact number of sex chromosomes operating in this species is yet to be determined. Irrespective of the exact location on the chromosomes, our new marker system enables accurate identification of the genetic sex in the common toad. While the majority of amphibian species to which genetic sexing methods have been established feature XX/XY sex determination (Alho et al., 2010; Baroiller & D’Cotta, 2016; Lambert et al., 2019; Nemesházi et al., 2020; Xu et al., 2021; Yoshimoto et al., 2008), our new marker set provides a cheap and easy-to-use method for future studies aiming to understand sex-reversal mechanisms in an anuran species with ZW/ZZ sex determination.

Our field study on several hundreds of adult toads found only a single case of sex reversal: 3 out of our 4 sex markers confirmed that the W chromosome was present in the DNA sample of one adult male captured at an agricultural site (the fourth marker gave inconclusive result). In the lack of further swab samples from this animal, we cannot completely exclude the possibility of contamination. Nevertheless, we had the highest number of phenotypic male samples from the pond where this individual was captured (Figure S1), thus, finding a single sex reversal at this site is compatible with the idea of an existing but very low frequency of sex reversal in the studied common toad populations. This almost complete lack of sex reversal is surprising, because we found many EDCs in the studied ponds, with higher concentrations in anthropogenic areas, as published in earlier papers (Bókony et al., 2021, 2018). Furthermore, we found a considerable number of female-to-male sex-reversed agile frogs *(Rana dalmatina)* in some of these ponds in the same years (Figure S1; Nemesházi *et al*., 2020). Thus, the lack of sex reversal in toads cannot be explained by the general lack of sex-reversing effects in the studied sites. Instead, this result may suggest that toad populations living in more polluted areas might have evolved resistance to sex reversal, thereby showing the same undisrupted sex development as their conspecifics in natural habitats. However, this interpretation is not supported by the results of our common garden experiment, because the effects of sex-reversing EDC treatments on the offspring of the studied toads did not depend on their original habitat type. Instead, they either all showed no sex reversal at low-concentration of both EDCs, or showed a 100% male-to-female sex reversal in the presence of high EE2 concentration (Figure 2). We only found a slight indication of habitat dependence of EDC susceptibility, suggesting that toads originating from anthropogenic habitats may be more, not less, susceptible to disrupted sex development: only urban toadlets displayed intersex gonads in a few cases when treated with ecologically realistic, low EDC concentrations. It remains to be tested if other EDC compounds or other concentrations within the range of the realistically low and close-to-maximum values that we applied here would reveal habitat-dependent sex-reversal probabilities in toads or any other species liable to sex reversal. Nevertheless, since most EDCs found in amphibian breeding habitats have estrogenic potential (Bókony et al., 2018), our treatments provide a good overall representation of the estrogenic EDC effects likely present in the field.

As a possible explanation for our results, survival rate of sex-reversed juveniles might be low in the wild, resulting in low prevalence of sex-revered individuals among adults. The environmental stimuli that cause sex reversal may have other developmental effects that might reduce survival; for example, heat stress in agile frogs increases both sex-reversal rate and mortality (Mikó et al., 2021). However, a meta-analysis found no significant relationship between chemically induced sex reversal and mortality in aquaculture fish (Alistair McNair Senior et al., 2012), and similarly, in our present study, toadlet survival was not reduced in the treatment group that showed 100% male-to-female sex reversal (Table S2). Moreover, sex-reversed individuals may show different behaviour (Li, Holleley, Elphick, Georges, & Shine, 2016; Alistair McNair Senior, Lokman, Closs, & Nakagawa, 2015). If their altered behaviour also affects their microhabitat use or results in changed activity during the breading season, these individuals might be harder to find by conventional capturing methods. There is currently no published information on the survival and behaviour of sex-reversed individuals in nature, so testing the above ideas will require further research.

As an alternative explanation that is not mutually exclusive with the above hypotheses, we speculate that the common toad may be relatively resistant to sex reversal, regardless of habitat type. In all other anuran species studied so far for sex reversal in free-living populations, female-to-male sex reversal was found in noticeable numbers (Alho et al., 2010; Lambert et al., 2019; Nemesházi et al., 2020; Xu et al., 2021), and a few cases of male-to-female sex reversal were also indicated (Lambert et al., 2019). There are several differences between the common toad and the previously studied anuran species, which might contribute to the apparent difference in sex-reversal frequencies found in their wild populations. First, all species studied so far belong to the family Ranidae, whereas the common toad is a member of Bufonidae; and different phylogenetic lineages may show different sensitivity for certain sex-reversing conditions (Chardard, Penrad-Mobayed, Chesnel, Pieau, & Dournon, 2004; Hayes, 1998; Orton & Tyler, 2015; Tamschick et al., 2016). Second, toads produce defensive toxins from cholesterol, the precursor of steroid hormones (Daly, 1995), and they have been selected for resisting autotoxicity (Moore, Halliday, Rowell, Robinson, & Keogh, 2009). This might have conferred them tolerance to other chemical perturbations which mimic the effects of steroid hormones (including estradiols and “stress hormones”), similarly to the cross-resistance provided by tolerance to certain pesticides in other anurans (Hua, Jones, & Relyea, 2014). Third, sex reversal may be triggered by not only chemical but also thermal stimuli, and different species may have adapted to different temperatures. If sex reversal in free-living amphibians occurs mostly due to extreme temperatures (Lambert et al., 2018; Mikó et al., 2021), the lack of sex reversal in common toads might be explained by their higher tolerance to heat. In line with this idea, the breeding season starts ca. one month later in spring for common toads than for agile frogs in our study region, and accordingly, we found female-to-male sex reversal in agile frogs (Nemesházi et al., 2020) but not in common toads among free-living adults, and we found the same difference between the two species in an experimental study of heat-induced sex reversal (Ujszegi et al., unpubl. data). Similarly, evolution of different temperature thresholds for sex reversal was suggested to explain the finding that in a reptile, *Pogona vitticeps*, sex reversal is absent in the hottest part of the species’ range (Castelli et al., 2021).

What makes different populations and species more or less susceptible to sex reversal is an important question for evolutionary ecology as well as for conservation biology. Possible reasons include constraints such as the degree of sex-chromosome heteromorphy (Miura, Ohtani, Ogata, & Ezaz, 2016) and direct or indirect selection pressures. For example, artificial selection for increased fecundity in females can indirectly affect male sensitivity to estrogenic disruption of testis development and spermatogenesis (Spearow, Doemeny, Sera, Leffler, & Barkley, 1999). Such selection pressures may force populations to evolve or plastically modulate any element involved in the biochemical pathway that translates environmental stimuli into sex (Castelli, Whiteley, Georges, & Holleley, 2020), e.g. by mutation of genes encoding hormone receptors (Castañeda Cortés, Arias Padilla, Langlois, Somoza, & Fernandino, 2019; Hamilton et al., 2020). Thus, the vulnerability of phenotypic sex development may be shaped by multiple forces, which might explain why researchers so far had mixed success in finding clear-cut relationships of sex-reversal rate with environmental factors such as climate (e.g. Castelli *et al*., 2021 vs. Dissanayake *et al*., 2021) and urbanization (Lambert *et al*., 2019 vs. Nemesházi *et al*., 2020) or with taxonomy (Alistair M Senior & Nakagawa, 2013). Even when a clear correlation is present, the underlying mechanisms are difficult to ascertain: for example, estrogenic pollution in river stretches is associated with high frequency of intersex in fish but not with polymorphisms in genes involved in responses to EDCs (Hamilton et al., 2020). Our present results with common toads add to this complex picture, emphasizing the need for further research on sex reversal in a wide diversity of species.

Building on our accumulated understanding from laboratory experiments on how environmental perturbations affect sex and from theoretical models on how sex reversal may impact population dynamics and evolution, the time is ripe for empirical studies on the causes and consequences of sex reversal in wild populations in the Anthropocene.

## Acknowledgments

We thank Viktória Verebélyi, Márk Szederkényi, and Patrik Katona for help with animal handling and data archiving, and all members of the Lendület Evolutionary Ecology Research Group for insightful discussions. We are grateful to Flóra Kerekes and Eszter Kása for preparing the histological sections, Beata Rozenblut-Kościsty for help with interpreting histological images, and Júlia Halász and Zsuzsanna György for help with cloning. DNA work was carried out in collaboration between the laboratories of Conservation Genetics Research Group of the University of Veterinary Medicine (Budapest; primer development and genetic sexing), the MTA-DE Lendület Evolutionary Phylogenomics Research Group (Debrecen; RADSeq) and the Institute of Genetics and Biotechnology, Hungarian University of Agriculture and Life Sciences (Budapest; cloning). The study was funded by the National Research, Development and Innovation Office of Hungary (NKFIH grants 115402 & 135016 to VB). VB was supported by the János Bolyai Research Scholarship of the Hungarian Academy of Sciences and by the New National Excellence Program of the Ministry for Innovation and Technology from the source of the National Research, Development and Innovation Fund (ÚNKP-20-5 & ÚNKP-21-5). EN was supported by the Ministry of Human Capacities (National Program for Talent of Hungary, NTP-NFTÖ 17-B-0317). EB was supported by the New National Excellence Program of the Ministry for Innovation and Technology (ÚNKP-21-2). None of the funding sources had any influence on the study design, collection, analysis, and interpretation of data, writing of the paper, or decision to submit it for publication.

## Data Accessibility Statement

Sequence data of the sex-linked markers are deposited to NCBI GenBank (accession numbers: OK507208-OK507215).

Individual-based metadata are stored in FigShare (DOI: 10.6084/m9.figshare.16809991).

## Author contributions

VB designed and supervised the study. Field work was done by BÜ and VB. Animals in the laboratory were raised by EN, NU, and VB; dissection and phenotypic sexing was performed by NU. RADseq was performed by GS, LL, and LS. All further DNA work was conducted by EN, EB, and NV, supervised by EN. EN and VB wrote the manuscript. All authors proofread the manuscript and gave final approval for publication.

## Notes

https://doi.org/10.6084/m9.figshare.16809991

### Competing Interest Statement

The authors have declared no competing interest.

