## Supplementary Material for "Novel genetic sex markers reveal unexpected lack of, and similar susceptibility to, sex reversal in free-living common toads in both natural and anthropogenic habitats"

**Table of contents**

1. Capture sites ........................................................................................ 2.
2. Experimental procedures........................................................................ 5.
3. DNA extraction...................................................................................... 10.
4. Details of sexing-primer design and sex-marker optimization............................... 11.
5. Genome BLAST of the sex markers ......................................................... 16.
Supplementary References ........................................................................ 17.

### *1. Capture sites*

Twelve of our 14 capture sites have been described in detail in an earlier paper (Bókony et al. 2018). These 12 sites were sampled in 2017 to investigate the chemical pollutants in breeding ponds and the toads' reproductive performance (Bókony et al. 2018), toxin production (Bókony et al. 2019), and sexual dichromatism (Ujhegyi and Bókony 2020). For the current study, we additionally captured adults at a natural site in 2016 (Garancsi-tó), and one pair at an urban site in 2017 which was not included in the previous papers due to the small sample size. The characteristics of each study site and the number of captured toads are given in Table S1. Pond locations are shown in Figure S1.

We categorized habitat type based on geoinformatics measurements of land use in a 500-m wide belt around each pond, as described in detail in an earlier paper (Bókony et al. 2018). We measured the area of 6 land-use categories (Table S1): “natural” vegetation (e.g. woodlands, non-agricultural meadows), arable fields, pastures, residential areas, public built areas (e.g. commercial and industrial areas), and roads with vehicular traffic. Using these 6 landscape variables we performed a principal component analysis (PCA), which yielded two axes with >1 eigenvalue, explaining 80.2% of variation in total; urban and agricultural landscape areas loaded on the first and second axis, respectively (loadings, PC1: arable fields -0.30, pastures -0.21, natural vegetation -0.27, residential areas 0.53, public, built areas: 0.46, roads: 0.55; PC2: arable fields 0.60, pastures 0.43, natural vegetation -0.66, residential areas 0.05, public, built areas: 0.10, roads: 0.04). The 14 capture sites separated along these two PCA axes into three clusters (Figure S2), corresponding to our subjective categorization of habitat type except that one site (Merzse) was assigned between natural and agricultural areas (note, however, that this site was not used for the common garden experiment).

**Table S1. Geographical coordinates, land use characteristics, and sample sizes for the 14 capture sites in the present study.** Note that the numbers of phenotypic males and phenotypic females reflect capture success and not population sex ratio at each site.

| Pond (abbreviation) | Habitat type | Coordinates | | Proportion of landscape cover | | | | | | Adult toads (males, females) | Number of clutches in common garden |
|---|---|---|---|---|---|---|---|---|---|---|---|
| | | N° | E° | Arable fields | Pastures | Natural vegetation | Residential areas | Public, built areas | Roads | | |
| Anyácsapuszta (A) | agricultural | 47.582 | 18.697 | 0.802 | 0.051 | 0.145 | 0 | 0 | 0.007 | 15*, 6 | 2 |
| Bajdázó (B) | natural | 47.904 | 18.978 | 0 | 0.022 | 0.97 | 0 | 0 | 0.024 | 7, 7 | 7 |
| Erzsébet-ér (E) | urban | 47.429 | 19.134 | 0.015 | 0.102 | 0.370 | 0.324 | 0.124 | 0.063 | 1, 1 | 1 |
| Garancsi-tó (Ga) | natural | 47.624 | 18.806 | 0.002 | 0.056 | 0.859 | 0.066 | 0.001 | 0.015 | 26, 26 | 0 |
| Göd (Göd) | urban | 47.685 | 19.130 | 0 | 0 | 0.248 | 0.431 | 0.033 | 0.053 | 14, 12* | 11 |
| Gyermely (Gy) | agricultural | 47.613 | 18.652 | 0.735 | 0.014 | 0.213 | 0.001 | 0.027 | 0.011 | 17*, 0 | 0 |
| Határrét (Ha) | agricultural | 47.645 | 18.911 | 0.484 | 0.137 | 0.284 | 0.07 | 0 | 0.026 | 27, 12 | 11 |
| János-tó (J) | natural | 47.714 | 19.020 | 0 | 0 | 0.987 | 0 | 0 | 0.012 | 18, 16 | 14 |
| Merzse (M) | natural / agricultural | 47.446 | 19.284 | 0.341 | 0.068 | 0.584 | 0 | 0 | 0.011 | 9, 1 | 0 |
| Perőcsény (Pe) | agricultural | 47.986 | 18.841 | 0.346 | 0.141 | 0.498 | 0 | 0 | 0.014 | 12, 5 | 3 |
| Pesthidegkút (Ph) | urban | 47.569 | 18.955 | 0.013 | 0 | 0.156 | 0.724 | 0.031 | 0.077 | 22, 12 | 8 |
| Pilisvörösvár (Pv) | urban | 47.610 | 18.920 | 0.004 | 0.024 | 0.27 | 0.531 | 0.083 | 0.077 | 22, 12 | 7 |
| Pilisszentiván (Ps) | urban | 47.607 | 18.908 | 0 | 0 | 0.282 | 0.455 | 0.173 | 0.076 | 11, 11 | 10 |
| Szárazfarkas (Sz) | natural | 47.734 | 18.819 | 0 | 0 | 0.988 | 0 | 0 | 0.012 | 15, 15 | 15 |

*One individual from each of these 3 groups could not be sexed genetically.

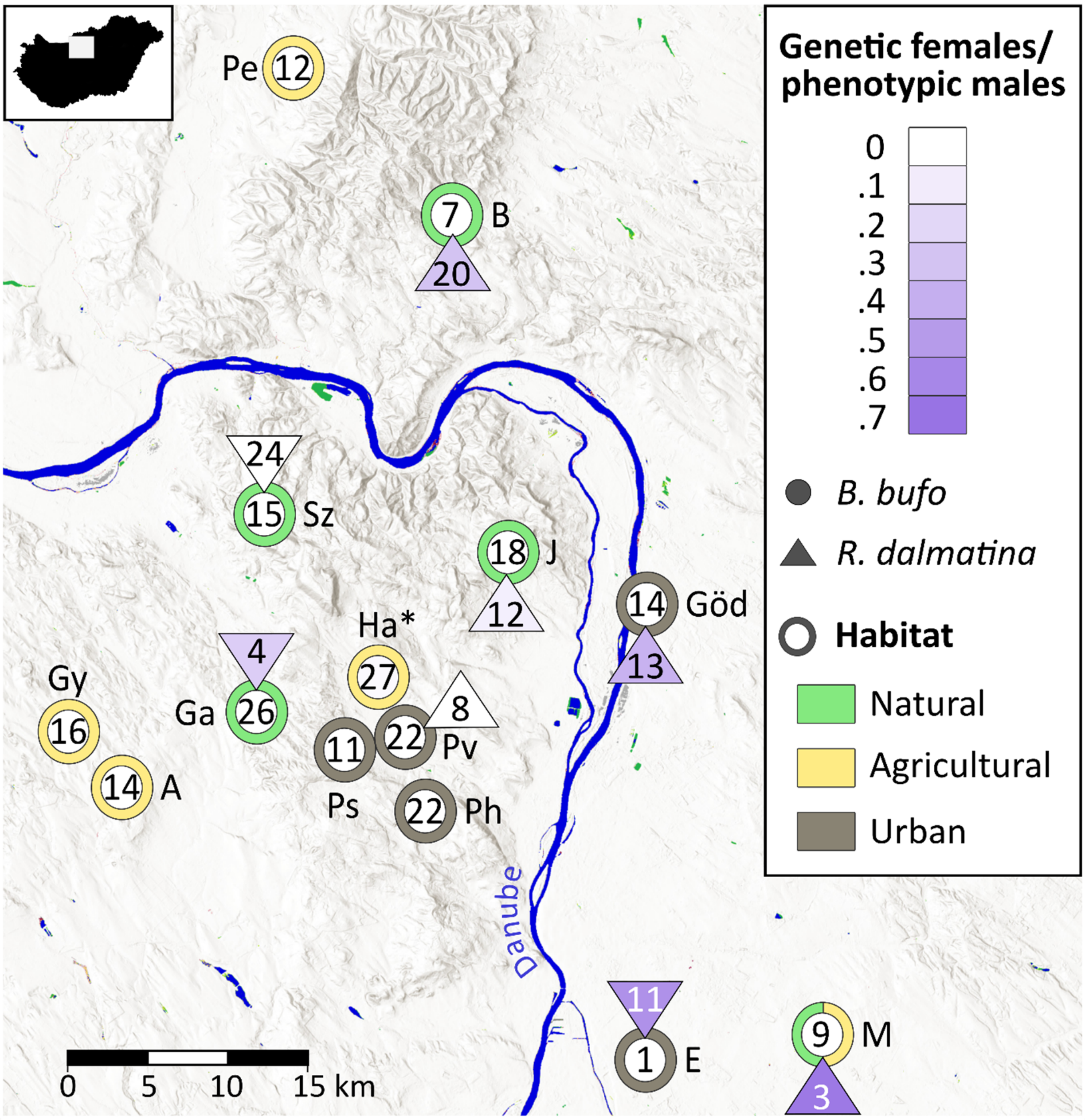


**Figure S1. The number of successfully genotyped wild-caught males in each breeding pond and the respective ratios of genetic females among phenotypic males.** Common toads *(Bufo bufo)* from the present study are shown with circles. For comparison, agile frog *(Rana dalmatina)* data from the same breeding ponds are shown with triangles: these data suggest that sex-reversing effects are present in many of these breeding sites at biologically relevant levels (source: Nemesházi et al. 2020). The only sex-reversed (ZW) male common toad was found in Határrét, marked by an asterisk (the proportion of genetic females in phenotypic males was 0.037 in this pond). Hillshade (ESRI World hillshade) and water (Global Surface Water Transitions) displays were downloaded using the QuickMapServices module in QGIS 3.18.3. Art work was done in Inkscape 0.92.

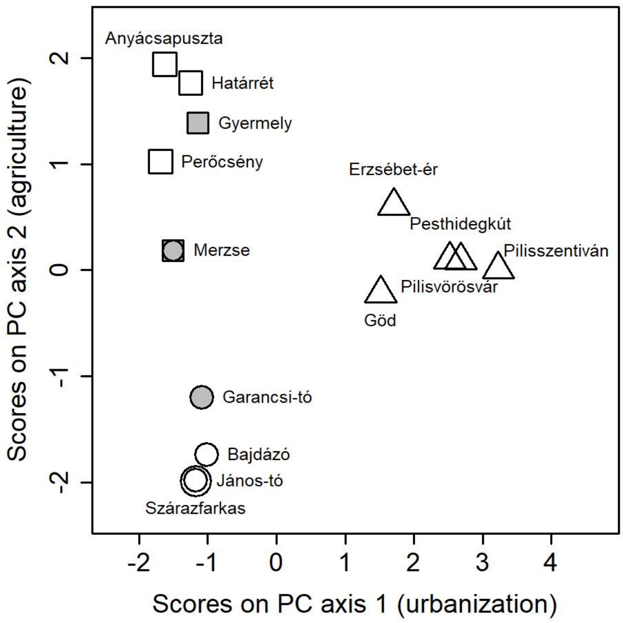


**Figure S2. Grouping of the capture sites of common toads along gradients of urban and agricultural land use.** Symbol shape indicates habitat type (circles: natural, squares: agricultural, triangles: urban; a site intermediate between natural and agricultural habitats is shown with a circle inside a square, and two overlapping natural sites are shown with two concentric circles). Symbol colour indicates whether the site was used only for sampling adults (grey) or also for the common garden experiment (white).

### *2. Experimental procedures*

Adult toads were captured by a drift fence with pitfall traps in 2016 and by hand in 2017. Whenever we captured gravid females at a site in 2017, all toads were transported to our laboratory for the common garden experiment, where temperature was 20 ± 1.55 °C and artificial light-dark cycles mimicked the natural photoperiod. We housed each pair in a 52 × 37 × 33 cm plastic box filled with 15 L reconstituted soft water (RSW; 48 mg $NaHCO_3$, 30 mg $CaSO_4 \times 2 H_2O$, 61 mg $MgSO_4 \times 7 H_2O$, 2 mg KCl added to 1 L reverse-osmosis filtered, UV-sterilized tap water) and containing 4 vertical wooden sticks as spawning substrates. Each box housed one male and one female haphazardly chosen from the individuals captured at the same pond. The pairs were allowed to spawn for one week (they spawned 0-7 days after capture, with a median of 2 days; 87% spawned within 3 days), after which they were released along with most of their eggs at the pond where they had been captured.

From each pair, we kept ca. 30 eggs in the lab until hatching. When the embryos became free-swimming tadpoles, we selected 6 healthy-looking individuals (i.e. no visible deformities or abnormal behaviour) from each family and moved each tadpole into a 2-L plastic box filled with 1 L RSW treated according to the chemical treatment group assigned to each individual (as detailed in the main text). The remaining tadpoles were released to the pond where their parents had been captured. We raised the tadpoles to metamorphosis by feeding them with chopped commercial spinach *ad libitum* and changing their rearing water twice a week. We applied the treatments throughout the entire larval development because we are not aware of any information about the timing of a sensitive period of sex determination in common toads. When a tadpole started metamorphosis (i.e. appearance of forelimbs), we decreased the water level to 0.1 L and slightly tilted the container to allow the animal to leave the water. When it completed metamorphosis (i.e. disappearance of the tail), we moved it into a clean rearing box containing wet paper towels as substrate and a piece of egg carton as shelter, which were changed every two weeks. We fed the toadlets *ad libitum* with springtails and small crickets, amended with a 3:1 mixture of $CaCO_3$ and Promotor 43 powder (Laboratorios Calier S.A., Barcelona, Spain) containing vitamins and amino acids. We raised the juveniles for 119-178 (median: 160) days after completion of metamorphosis; dissections took place between October 6 and November 10, 2017. By this time, our initial sample size of 534 decreased by 117 because two toadlets escaped from their boxes and 115 individuals died, most of them (N=78) in the 3 mg/L glyphosate treatment (Table S2); this high level of mortality was unexpected based on our earlier study on common toad tadpoles from the same region (Mikó et al. 2017). Additionally, one female that died shortly before the start of dissections was also dissected and sexed. The timing of dissection was balanced among the animals from the three habitat types such that natural, agricultural and urban individuals were systematically rotated during the one-month period. We euthanized the toadlets by a one-hour immersion into a room-temperature water bath of 5.4 g/L MS-222 (Sigma E10521) buffered to neutral pH with the same amount of $Na_2HPO_4$, and we inspected the gonads under a stereomicroscope with 1-3× optical zoom (Figure S3 - Figure S4. All captures and experimental procedures were carried out according to the permits issued by the Government Agency of Pest County (Department of Environmental Protection and Nature Conservation) and the Budapest Metropolitan Municipality (Department of City Administration, FPH061/2472-4/2017). The experiments were further approved by the Ethical Commission of the Plant Protection Institute (ATK NÖVI).

**Table S2. Number of laboratory-raised individuals in each treatment.**

| Groups | ZW female | ZZ male | ZZ female | ZZ uncertain | Died | Total |
|---|---|---|---|---|---|---|
| **Control** | **36** | **46** | | **1** | **6** | **89** |
| Natural | 15 | 19 | | 1 | 1 | 36 |
| Agricultural | 8 | 7 | | | 1 | 16 |
| Urban | 13 | 20 | | | 4 | 37 |
| **Glyphosate 3 µg/L** | **35** | **39** | | **3** | **12** | **89** |
| Natural | 10 | 22 | | | 4 | 36 |
| Agricultural | 9 | 5 | | | 2 | 16 |
| Urban | 16 | 12 | | 3 | 6 | 37 |
| **Glyphosate 3 mg/L** | **7** | **4** | | | **78** | **89** |
| Natural | 1 | 1 | | | 34 | 36 |
| Agricultural | 4 | | | | 12 | 16 |
| Urban | 2 | 3 | | | 32 | 37 |
| **Solvent control** | **49** | **29** | | **1** | **8 (+2*)** | **89** |
| Natural | 16 | 14 | | | 4 (+2*) | 36 |
| Agricultural | 10 | 5 | | | 1 | 16 |
| Urban | 23 | 10 | | 1 | 3 | 37 |
| **EE2 1 ng/L** | **48** | **35** | | **1** | **5** | **89** |
| Natural | 22 | 12 | | | 2 | 36 |
| Agricultural | 8 | 7 | | | 1 | 16 |
| Urban | 18 | 16 | | 1 | 2 | 37 |
| **EE2 1 µg/L** | **44** | | **39** | | **6** | **89** |
| Natural | 14 | | 20 | | 2 | 36 |
| Agricultural | 11 | | 5 | | | 16 |
| Urban | 19 | | 14 | | 4 | 37 |
| ***Total*** | ***219*** | ***153*** | ***39*** | ***6*** | ***117*** | ***534*** |

* Two toadlets escaped their rearing boxes

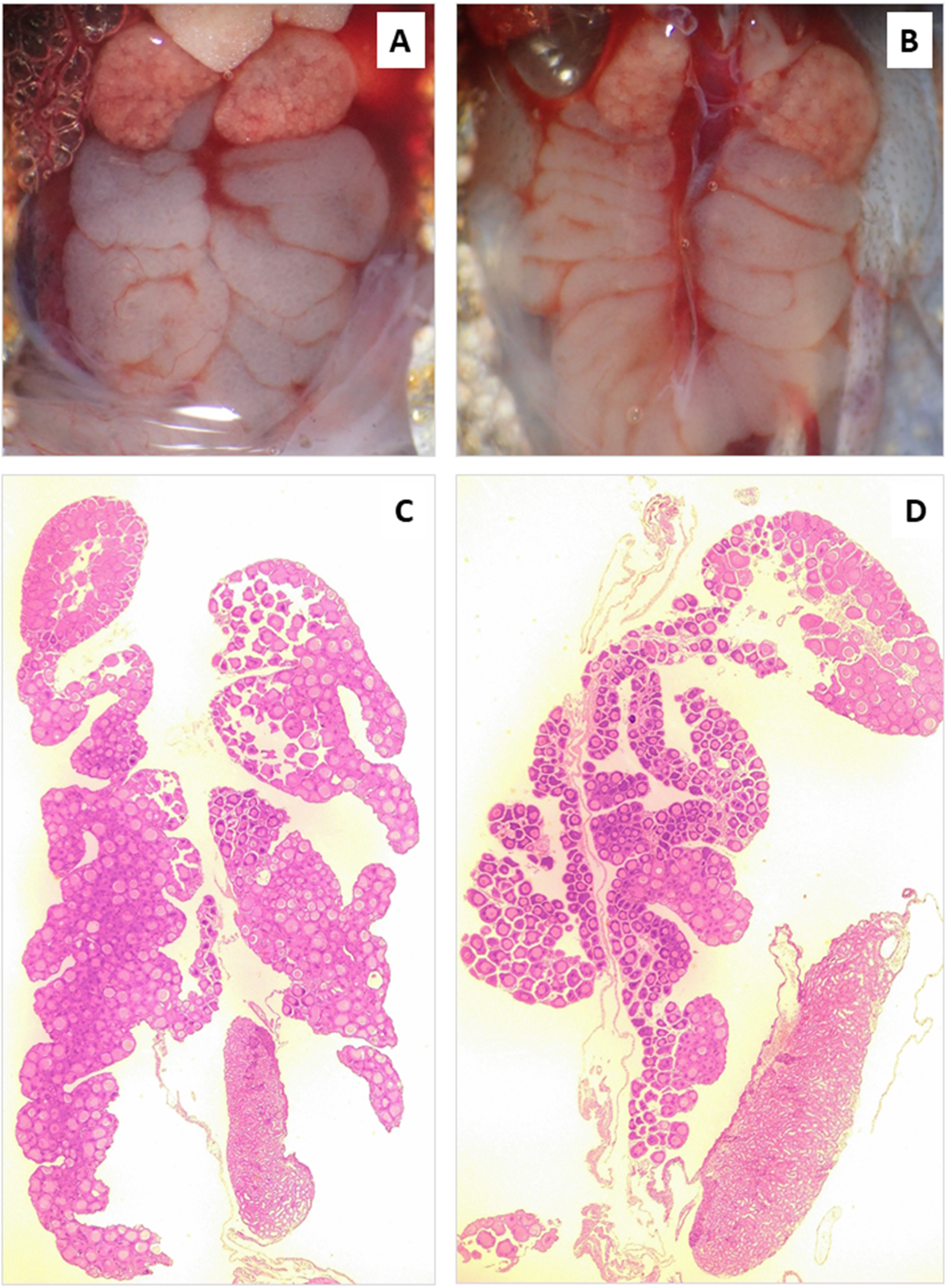


**Figure S3: Ovaries in juvenile common toads raised in the 1 µg/L EE2 treatment group.** Gross anatomy (A, B) and histology (C, D) of a concordant female (A, C) and a male-to-female sex-reversed individual (B, D).

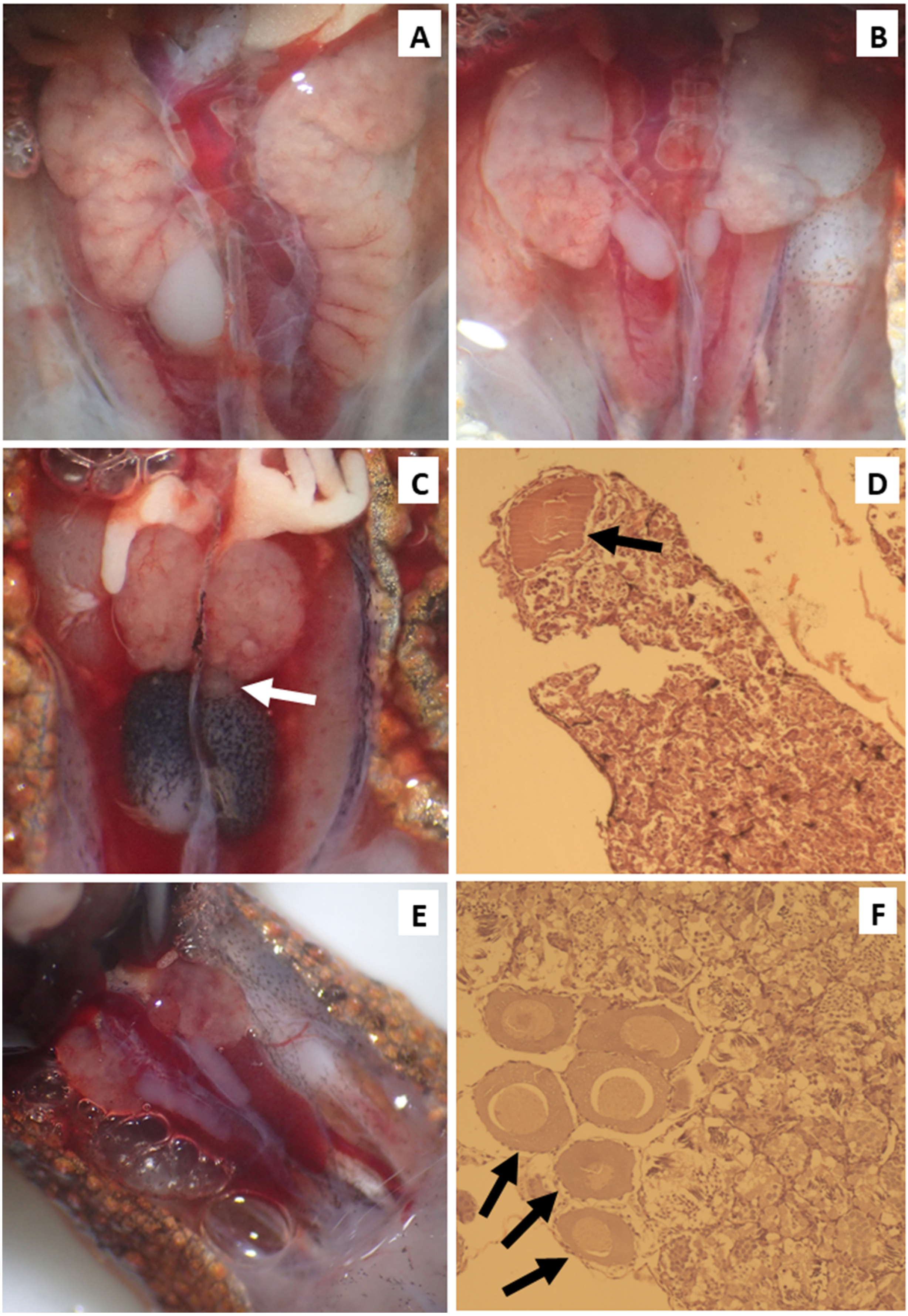


**Figure S4. Ambiguous gonads in juvenile common toads** (see next page for more info).

A) The gonad on the left is a normal testis and the gonad on the right is an ovary. This individual originated from an urban pond (Pesthidegkút) and was treated with 1 ng/L EE2.

B) The testes are abnormally shaped and the Bidder's organ on the right has an ovary-like structure. This individual originated from an urban pond (Pilisvörösvár) and was treated with 3 µg/L glyphosate.

C) A small ovary-like structure (white arrow) between the testis and Bidder's organ. Two individuals treated with 3 µg/L glyphosate had this morphology; both originated from urban ponds (Göd, Pesthidegkút).

D) Histological image of the gonad from one of the individuals showing the gross anatomy in panel C. A single oogonium (black arrow) is found in the testicular tissue.

E) Abnormal gross anatomy of testes and Bidder's organs in an individual from the solvent control group, originating from an urban pond (Pilisszentiván). The histological section of this individual was lost, so its phenotypic sex was categorized as uncertain.

F) Histological image of the gonad from the other individual whose phenotypic sex was categorized as uncertain. The cells shown by arrows may be testicular oogonia, or may belong to the Bidder's organ. Gross anatomy showed normal testes. This individual originated from a natural pond (János-tó) and was raised in the control group.

#### *3. DNA extraction*

For RADseq, we extracted genomic DNA from foot clips of 24 toadlets following the protocol of (Cserkész et al. 2015). Briefly, tissue samples were placed in 150 µl lysis buffer containing 0.2 % SDS, 100 mM Tris-HCl (pH = 7.5), 200 mM NaCl, 5 mM EDTA (pH = 8.0). To ensure effective digestion we added 20 µl Proteinase-K (20 mg/mL) (Thermo Scientific, MA, USA) to the mixture and incubated the samples overnight.at 55 °C. To remove RNA content, we added 5 µl (1 mg/mL) (Thermo Scientific, MA, USA) and incubated the tubes at room temperature (15 - 25 °C) for 15 minutes. Then, genomic DNA was washed with 0.5 V of ammonium-acetate (7.5 M) and an equal volume of chloroform-isoamylalcohol (24:1). The supernatant was transferred to a clean tube and DNA was precipitated with an equal volume of isopropanol and pelletised by heavy centrifuging. The pellets were washed twice with 70 % ethanol and resuspended in Tris-HCl (pH = 7.5) buffer. Isolates were checked for the presence of high molecular weight DNA using gel electrophoresis on 1% agarose-gel.

From buccal swab samples of adult toads collected in 2017, genomic DNA was extracted with either Bio-Tek Omega E.Z.N.A. Forensic DNA kit (99 individuals) or Quiagen QIAamp Investigator kit (200 individuals) following the manufacturers' instructions, except for the following modification. When using the E.Z.N.A. Forensic kit, we added 200 µl TL Buffer, 25 µl Proteinase K and 225 µl BL Buffer to each swab and incubated it at 60°C for 1 hour. From samples of adults captured in 2016, DNA was extracted with Thermo Fisher Scientific Geneaid Genomic DNA Extraction Kit (23 toe clips and one muscle sample from an animal found dead), and Bio-Tek Omega E.Z.N.A Tissue DNA kit (29 toe clips) following the original protocols with some modifications (at least 2-hr digestion followed by 30-min lysis). In the end of the DNA extraction with either one of the kits, we added 100 µl elution buffer to the columns containing the purified samples.

For all analyses of toadlets after RADseq, we extracted genomic DNA from foot clips by either Thermo Fisher Scientific Geneaid Genomic DNA Extraction Kit (54 juveniles from the control group) or Bio-Tek Omega E.Z.N.A Tissue DNA kit (363 juveniles including individuals from the control as well as the treated groups) as described above.

#### *4. Details of sexing-primer design and sex-marker optimization*

For c16, considerable size difference (in total 43 bp) resulting from InDels allowed unambiguous Sanger sequencing of both the Z and W allele. Sequences obtained from c2, c5 and c12 were unambiguous in the male and partially also in the three females, but double peaks appeared after a certain point in the latter sex. We suspected that this issue was caused by InDels between the respective W and Z copies of these loci and decided to untangle the noisy sequences manually by comparing alternative bases to the Z sequence obtained from the male (Figure S5). This process resulted in a candidate W sequence for each locus. Based on these candidate sequences, we designed W-specific PCR primers to enable separate sequencing of the respective W alleles (Table S3). We sequenced each locus in a total of 5 males and 5 females as follows: the original primers binding to both Z and W were used for sequencing males (i.e. product including the whole target sequence), while females were sequenced using either a forward or reverse W-specific primer. Alignment of the respective sequences confirmed the presence of InDels between the Z and W copies in all three loci.

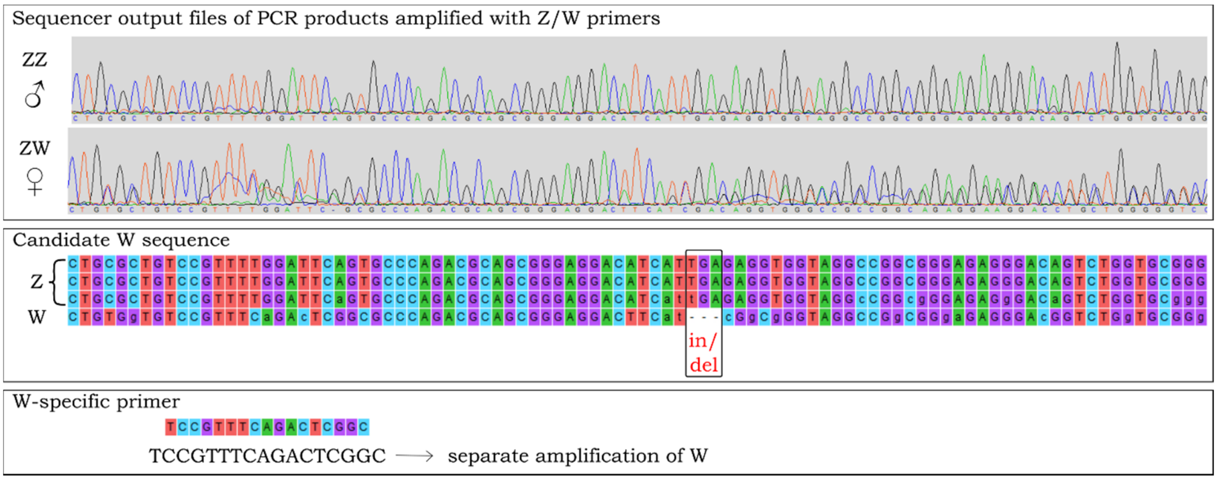


**Figure S5. Illustration of the designing process of W-specific primers based on sequences produced with primers that bind to both Z and W.** Respective sequences are shown below each other across all panels.

The InDel on c5 resulted in a 30 bp size difference between the Z and W alleles, which is suitable for detection on agarose gel, so first we designed sexing primers that embraced the sex-linked InDel (primers BbS5-F2 and Bb_c5-R in Table 1). However, after gel electrophoresis of the PCR products, the W-band was very faint in some of the females. We decided to combine this Z/W universal primer pair with the W-specific reverse primer (Bb_c5-W-R; see Table 1) designed earlier for sequencing, to obtain one (Z-linked) band in males and three (one Z- and two W-linked) bands in females (Figure S6). This primer combination amplified suitably bright Z and W bands in all individuals where the faint-W-product problem occurred earlier.

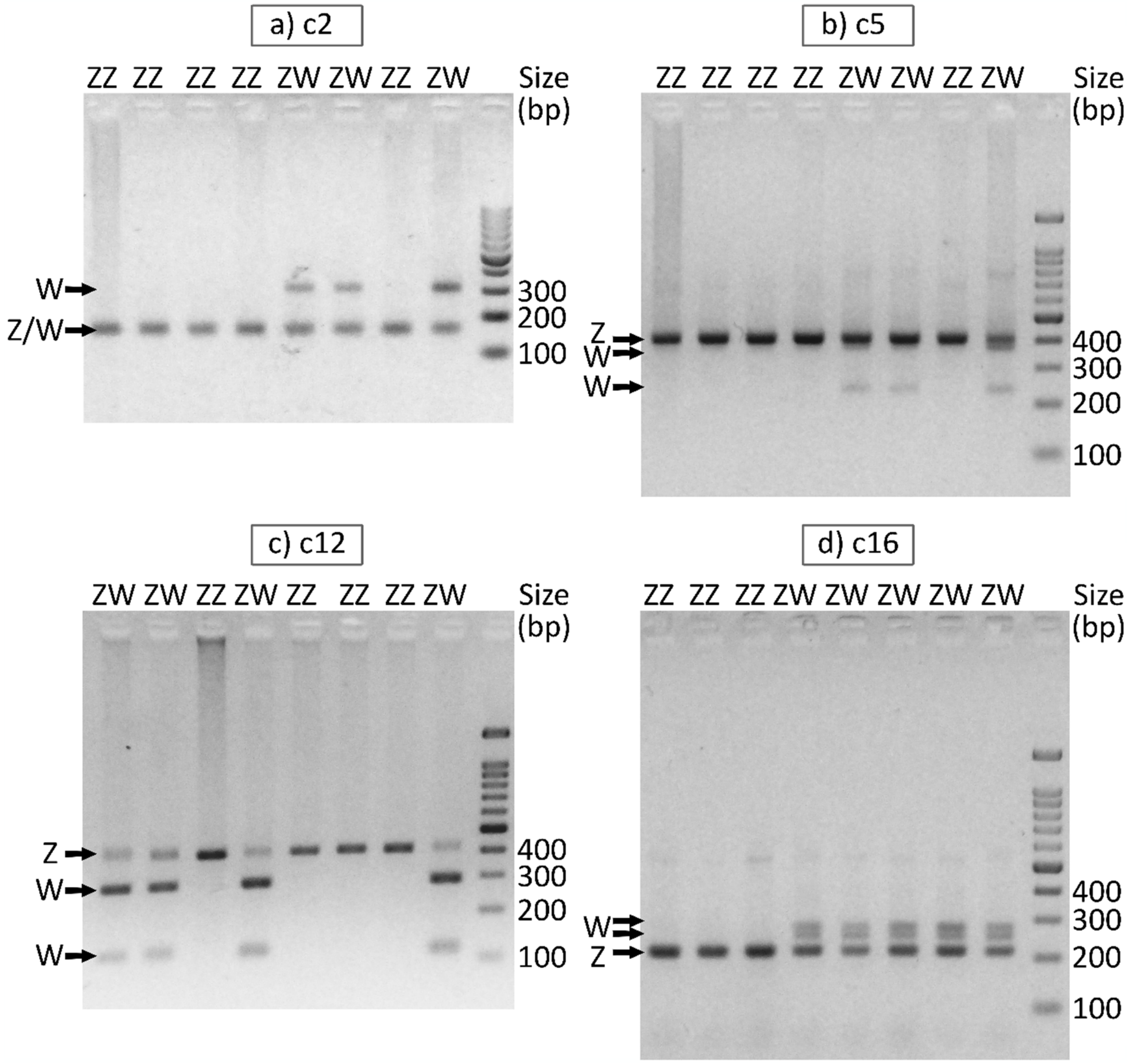


**Figure S6. Sexual genotypes identified on 2% agarose gel with each sex marker.** The primers used for each sexing PCR are listed in Table 1. Note that PCR products of c12 (panel c) were cut with TaiI enzyme before gel electrophoresis.

Size difference between the Z and W alleles of loci c2 and c12 was too short for detection on agarose gel (1 and 3 bp, respectively). However, consistent SNP differences between the Z and W alleles provided W-linked recognition sites for restriction enzymes. Therefore, PCR products of the sexing primers designed for these loci were digested enzymatically. While the PCR product of c12 was digested as described in the main text (see also Table 1 and Figure S6), c2 was digested by BseDI (10 U/µl, Thermo Scientific) in a two-hour digestion step was performed at 55°C, followed by heat inactivation at 80°C. Subsequent agarose gel electrophoresis resulted in one band in males (ZZ) and three bands in females (ZW; the W product was digested into two fragments, while the Z product remained intact) for both c2 and c12. Using c2, however, the W-band was missing or very faint in some females after multiple attempts of sexing. We obtained W sequences from 5 such females and found no discrepancies compared to other females: both the recognition site of the restriction enzyme and the binding site of the forward primer used for sexing matched across all the sequenced females. Because we could not identify the cause of insufficient W-specific digestion, we decided to amplify the c2 sex marker using the W-specific forward primer (previously used for sequencing W) together with the Z/W primer pair instead of digesting the PCR products (Table 1 and Figure S6). We could not exclude the possibility that the insufficient performance of the above-described restriction method was caused by mismatches present at the reverse primer's binding site. However, successful amplification of the W product after the introduction of a second forward (but not another reverse) primer suggests that the reverse primer's binding success to W was not a major issue here.

Because the RAD sequence of c16 suggested the presence of many SNPs across the locus, the forward primer contained a supposedly W-specific nucleotide at a distance of 3 positions from the 3' end of the primer. As a result, the Bb_c16-F2 + Bb_c16-R2 primer pair (Table S3) amplified W more efficiently compared to Z and it produced a single band in each sex (W in ZW females and Z in ZZ males), but the band size differed between sexes. The obtained sequences revealed that there is overall 42 bp size difference between the Z and W copies due to the presence of a total of three InDels. We designed further primer pairs that would bind to Z and W with the same efficiency (including a total of 40-43 bp InDel difference), which all produced a surprising sex-linked pattern: one band in males and three bands in females (see Table 1, Table S3 and Figure S6). We attempted to obtain sequence of the unexpected W band by cloning the respective PCR products with both the BbS16-F3 & BbS16-R and the Bb_c16-F & Bb_c16-R2 primer pairs. However, after purification, the sample always contained some of the shorter fragments as well, and the plasmid vector integrated those instead of the product of interest.

**Table S3. Primers used for Sanger sequencing of potentially sex-linked loci identified by RADSeq.**

| Locus | Primer | Primer sequence (5'->3')* | Target | Product (bp) | Annealing (°C) | Comment |
|---|---|---|---|---|---|---|
| c1 | Bb_c1-F2 | ACTTTGTGTGCTTAGTGTCCG | Z & W | 185 | 65 [1] | Ambiguous sequence |
| | Bb_c1-R2 | GGTTCCTACCTTGACACCCTA | | | | |
| c2 | Bb_c2-F | AGGACCTGTGTGGTCTGT | Z & W | Z: 385 | 68-64 [2] | Good in ZZ males |
| | Bb_c2-R | ATCATCGAAGGGAAGAGCCG | | W: 386 | | |
| | Bb_c2-W-F | TGTTCTATGCACTATGTGG | W | 311 | 65-57 [2] | |
| | Bb_c2-R2 | CGAAGGGAAGAGCCGTC | | | | |
| c3 | Bb_c3-F | GCAGGAGACTACCCACGGA | Z & W | 335 | 57 [1] | No sex-linked pattern |
| | Bb_c3-R | AGATCGCCGCTCAGAAGGTCG | | | | |
| c5 | Bb_c5-F | AGGATGACTGGCTTGATC | Z & W | Z: 477 | 61 [1] | Good in ZZ males |
| | Bb_c5-R | CTGGACGTATGTTCTCCACG | | W: 447 | | |
| | Bb_c5-W-F | CATCCGATGTTCCCAGCAGT | W | 346 | 68-64 [2] | |
| | Bb_c5-R | CTGGACGTATGTTCTCCACG | | | | |
| | Bb_c5-F | AGGATGACTGGCTTGATC | W | 339 | 68-64 [2] | |
| | Bb_c5-W-R | GGGCCAATTTTTTGGAGAAG | | | | |
| c9 | Bb_c9-F | TGCAGGGACCAAGCTAATCA | Z & W | 346 | 60 [1] | Ambiguous sequence |
| | Bb_c9-R | TGCACACCTTTGCATCTTCTG | | | | |
| c12 | Bb_c12-F | GTTCGGTCCCTCCTGAACG | Z & W | Z: 410 | 63 [1] | Good in ZZ males |
| | Bb_c12-R | CTCCTCAGGCCTAACCCGAT | | W: 407 | | |
| | Bb_c12-W-F | TCCGTTTCAGACTCGGC | W | 309 | 68-64 [2] | |
| | Bb_c12-R | CTCCTCAGGCCTAACCCGAT | | | | |
| c13 | Bb_c13-F | GTGGTTTCGGTCTTTCTTCCAG | Z & W | 422 | 68-64 [2] | No sex-linked pattern |
| | Bb_c13-R | CCCGTTTGTACACCTCTGTAT | | | | |
| c16 | Bb_c16-F | AGGTGGTTTCCATAGCGCTTTTA | Z & W | Z: 367 | 70-65 [3] | Good in ZZ males |
| | Bb_c16-R2 | AATGTCAGATGCGGGTCGG | | W: 410 & ca. 450 | | |
| | Bb_c16-F2 | TATGGAGCCTTAAAGGGGTGG | Z & W | Z: 313 | 70-65 [3] | If W is present, it is amplified predominantly |
| | Bb_c16-R2 | AATGTCAGATGCGGGTCGG | | W: 355 | | |
| c17 | Bb_c17-F | TGAGAACGTTTATGCCTCGCT | Z & W | 458 | 58 [1] | No sex-linked pattern |
| | Bb_c17-R | CACCGATGCCCAACCACTTA | | | | |

[*] Nucleotides highlighted in colour are SNP positions, and are either the W (red) or the Z (blue) versions. These primers bind to both sex chromosomes nevertheless.

[1] PCRs were performed as follows: 94°C for 2 min, 35 cycles of 94°C for 30 sec, annealing (see column "Annealing") for 30 sec, and 72°C for 60 sec, followed by 72°C for 10 min.

[2] The above PCR settings were modified so during first 13 cycles the annealing temperature decreased gradually between the temperatures indicated in column "Annealing" and the remaining cycles were performed with the lowest annealing temperature.

[3] The touch-down period lasted for the first 10 cycles.

#### *5. Genome BLAST of the sex markers*

Whole genome sequence of the common toad was recently published in the NCBI Genome database by the Wellcome Sanger Institute (assembly aBufBuf1.1). We performed genome blast with each of our 4 sex-linked loci to determine which chromosome they are located on. Nucleotide BLAST was performed using the Megablast algorithm on the NCBI website (https://blast.ncbi.nlm.nih.gov; word size: 16; without masking for low complexity regions). We downloaded the hit table for both the W and Z alleles of each locus and sorted the hits first by E-value and second by the percentage of identity. We accepted the best hit if its E-value was at least 5 orders of magnitude lower compared to the second best hit (the same threshold was used in Nemesházi et al., 2020). We also checked if the query cover contained the whole length of the known allele sequence (the latter ranged between 367 bp for the c16 Z allele and 477 bp for the c5 Z allele).

Based on the NCBI genome blast, we could identify three out of the four sex markers (c2, c12 and c16). The query cover was 100% in all of these cases and for each marker the best hit based on the Z allele was the same locus as the best hit based on the W allele. The percentage of identity was similar or higher for the Z sequences (98.5-99.5%) compared to the W sequences (87.6-97.9%), indicating that the W chromosome might be missing from the available genome assembly. The genome-assembly information indicated that c12 was located on chromosome 5 (accession LR991671.1; the c12 Z allele showing the overall highest, 99.5% identity), while c2 and c16 were localized on two different, unplaced scaffolds (c2 on CAJIMN010001189.1 and c16 on CAJIMN010000039.1). The full sequence report available for the genome assembly (last updated on 2021.01.28) showed that the scaffold containing c16 is currently suggested to belong to chromosome 6, while there was no indication of the chromosome where the other scaffold might belong.

Genome blast showed that highly similar sequences to locus c5 were present on several different chromosomes (including multiple locations on chromosome 5 and others) with 95.8-98.11% query cover: we found sequences with 98.7% identity with the Z allele and maximum 95.89% identity with the W allele. However, primer blast indicated that the c5 PCR primer pairs that we used for either sequencing or sexing would not likely amplify any of these loci (i.e. minimum three mismatches were present between one or both primers and their potential binding sites in case of each primer pair used). Indeed, the touch-down PCR protocol that we used for genetic sexing (see Methods) resulted in a sex-linked product pattern suggesting that, although the location of c5 within the genome is unclear, the sexing primers amplified fragments from the sex chromosomes (but at least the female-specific fragments were surely amplified from the W chromosome, because these were detected only in females).
